# Advancing mRNA subcellular localization prediction with graph neural network and RNA structure

**DOI:** 10.1101/2023.12.14.571762

**Authors:** Fuyi Li, Yue Bi, Xudong Guo, Xiaolan Tan, Cong Wang, Shirui Pan

## Abstract

**Motivation:** The asymmetrical distribution of expressed mRNAs tightly controls the precise synthesis of proteins within human cells. This non-uniform distribution, a cornerstone of developmental biology, plays a pivotal role in numerous cellular processes. To advance our comprehension of gene regulatory networks, it is essential to develop computational tools for accurately identifying the subcellular localizations of mRNAs. However, considering multi-localization phenomena remains limited in existing approaches, with none considering the influence of RNA’s secondary structure.

**Results:** In this study, we propose Allocator, a multi-view parallel deep learning framework that seamlessly integrates the RNA sequence-level and structure-level information, enhancing the prediction of mRNA multi-localization. The Allocator models equip four efficient feature extractors, each designed to handle different inputs. Two are tailored for sequence-based inputs, incorporating multilayer perceptron and multi-head self-attention mechanisms. The other two are specialized in processing structure-based inputs, employing graph neural networks. Benchmarking results underscore Allocator’s superiority over state-of-the-art methods, showcasing its strength in revealing intricate localization associations.

**Availability:** The webserver of Allocator is available at http://Allocator.unimelb-biotools.cloud.edu.au; the source code and datasets are available at https://github.com/lifuyi774/Allocator

## Introduction

The asymmetric distribution of mRNA has been discovered to be closely associated with various cellular processes, laying the groundwork for local protein translation and organism development (Ephrussi et al., 1991; Weatheritt et al., 2014). This significance of asymmetric distribution is universal across different species (Long et al., 1997; Kloc et al., 2002). In particular, Bouvrette *et al*. found that around 80% of cellular RNA species exhibit non-uniform distribution, with these patterns being broadly conserved throughout evolution (Benoit Bouvrette et al., 2018; Li et al., 2021). Recent advancements in high-resolution imaging technologies have further revealed that mRNA asymmetrical distribution, known as mRNA subcellular localization, plays a crucial role in temporal and spatial control of gene expression (Martin & Ephrussi, 2009). For example, in the *D. melanogaster* embryo, *cytoplasmic Nos* mRNA binds to RBP Smaug, inhibiting its translation and triggering decay. However, in later development, *Nos* mRNA is localized to the posterior pole and replaces Smaug with RBP Oskar, thereby protecting transcripts from degradation. However, the extensive time and substantial financial resources required for wet lab experiments pose significant limitations on conducting thorough investigations, thus impeding our comprehensive understanding of the RNA localization mechanism.

In recent years, there have been various computational approaches for predicting mRNA subcellular localizations. They offer effective and time-saving alternatives to traditional laboratory experiments. These approaches can be categorized into two types. The first type regards localization prediction as a single-label multi-class task, where the goal is to identify a single localization for a given transcript. Models such as RNATracker (Yan et al., 2019), iLoc-mRNA (Zhang et al., 2021), mRNALoc (Garg et al., 2020), and SubLocEP (Li et al., 2021) adopted this strategy to build their predictive models. However, not all transcripts serve a single function; they may be localized in different compartments to fulfil different roles, as seen with *Nos* mRNA. Therefore, developing a single-label model falls short of providing comprehensive predictions for multiple localizations and fails to reveal the complex relationships between labels. It’s essential to consider the broad spectrum of multi-localization in model development. To address this issue, DM3Loc (Wang et al., 2021) and Clarion (Bi et al., 2022) were introduced by treating it as a multi-label task. DM3Loc leveraged a multi-head self-attention mechanism for predictions, while Clarion employed XGBoost based on a problem transformation strategy for classification. Both offer predictions for multiple labels simultaneously, advancing the state-of-the-art in mRNA subcellular localization prediction.

Nevertheless, DM3Loc and clarion are developed based on sequence information. In addition to the primary mRNA sequence, the secondary structure also plays a pivotal role in the localization process. Indeed, mRNAs are often targeted and bound by RNA-binding proteins (RBPs) via the *cis*-localization element, which can act as signals for delivery to a specific cellular localization (Das et al., 2021). For instance, in the 3’UTR of fruit fly *bicoid* mRNA, several 50-nt sequences, known as bicoid localization elements, have been identified. These elements form stem-loop structures that facilitate intermolecular interactions (Buxbaum et al., 2015). This insight has motivated us to incorporate mRNA secondary structure into the model construction. The similarity between RNA structure and graph deep learning enables the utilization of graph neural networks (GNNs) in this task. GNNs have proven effective in capturing intricate patterns and correlations, showcasing remarkable performance across diverse bioinformatics tasks (Réau et al., 2023; Kang et al., 2022). Recent years have witnessed the emergence of several influential GNN variants, each offering a distinct set of strengths: MPNN (message-passing neural network) passes information between nodes to update node representations; GCN (graph convolutional network) uses graph convolution operations for local information propagation; GIN (graph isomorphism network) aims to identify structural similarities and solve isomorphism challenges; GAT (graph attention network) applies attention to determine the weights of information exchange between different nodes.

In this study, we propose Allocator, a novel framework to leverage the power of graph deep learning and RNA secondary structures to advance the mRNA subcellular localization prediction. Allocator incorporates various networks in its architecture, including multilayer perceptron (MLP), self-attention, and graph isomorphism network (GIN). Allocator employs a parallel deep learning framework to learn two views of mRNA representations including sequence-based features and structural features. Then these learned features are combined and used to predict six subcellular localization categories of mRNA. Allocator has been demonstrated to be outstanding in revealing label correlations according to six multi-label evaluation metrics. To our knowledge, Allocator is the first approach that utilizes graph neural networks to capture mRNA secondary structure information to predict mRNA subcellular localization.

## Material and Methods

### Benchmark dataset

In this study, we curated a benchmark dataset comprising over 26,000 entries for mRNA subcellular localization in *Homo sapiens*. These entries were collected from RNALocate (Zhang et al., 2016) and DM3Loc (Wang et al., 2021). The dataset includes a total of 17,298 unique mRNA sequences in FASTA format. All sequences have undergone sequence similarity reduction using CD-HIT (Fu et al., 2012) with an 80% sequence identify threshold. Their lengths range from 186-nt to 30,609-nt, with an average length of 3689-nt. We then randomly divided this dataset into the training set, validation set, and testing set in an 8:1:1 ratio. **Table 1** displays the specific distribution for each subcellular localization within the three subsets. The ‘Sequences’ column represents the number of FASTA sequences, while other columns, such as ‘Nucleus’, count only the number of labels (localizations). It is clear that the label relationship between mRNA and localization labels is not limited to one-to-one but also one-to-many, which can be addressed as a multi-label problem setting. Furthermore, it is important to highlight our decision to utilize the original sequences rather than truncating sequences in order to preserve the relatively complete genomic information.

**Table 1.**
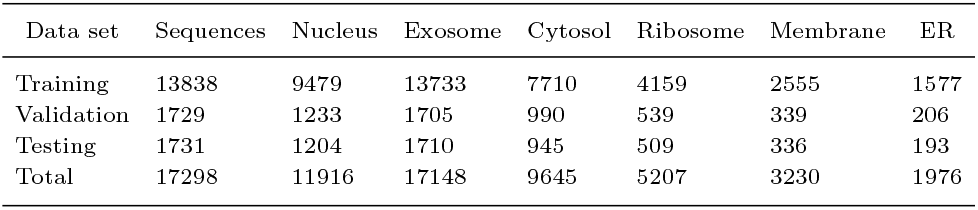
The distribution for six subcellular localizations in training, validation, and testing set.

### The framework of Allocator

Allocator is a multi-view parallel deep learning framework that is designed for mRNA multi-localization prediction. It is composed of three main modules: feature engineering, feature extractor, and prediction module, as depicted in **Figure 1**.

**Fig. 1.**
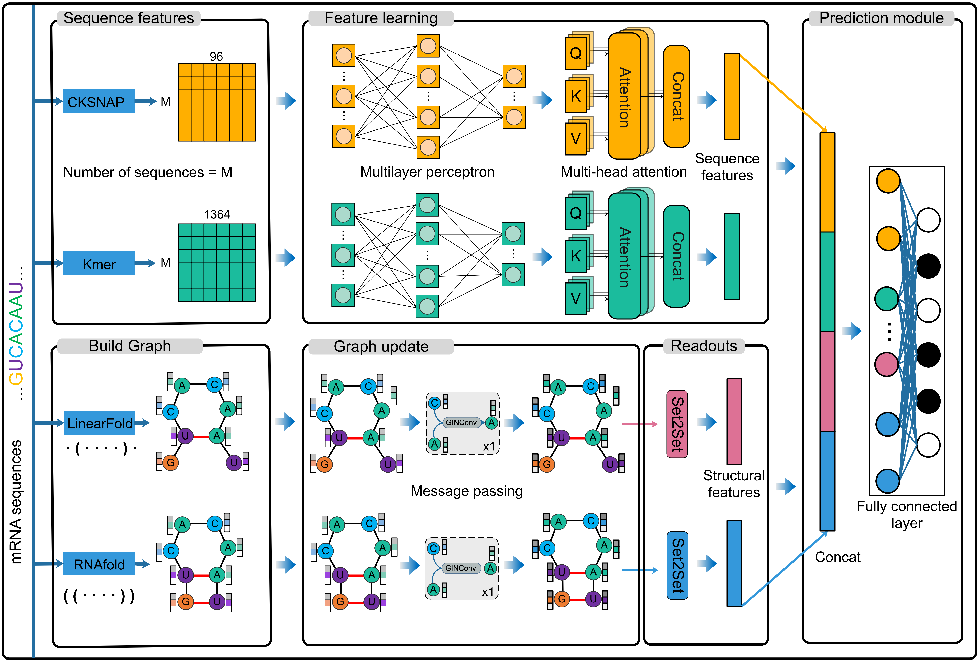
The deep learning framework of Allocator.

The feature engineering module involves preparing model inputs by leveraging information from both the RNA primary sequences and secondary structures. This step results in two sets of sequence context features and two sets of graph structural features, forming the foundation for the subsequent model training. Then, these features are processed through the feature extractor module to capture intricate and complex patterns. There are four parallel extractors, with two specifically for sequence context features and the other two for graph structure features. To ensure that both sequence context features and graph structural features have balanced dimensionality, we configured the four parallel extractors to each produce output features with a dimensionality of 128. Finally, the outcomes of these four feature extractors are concatenated and fed into the output module, consisting of two fully connected layers, to generate the predictive probabilities.

### Feature engineering

Regarding the initialise of sequence context features, we employed two encodings, *k*-mer and CKSNAP (*k*-spaced nucleic acid pairs), for extracting primary sequence characteristics (Chen et al., 2020; Lee et al., 2011). These encodings have proven their ability to capture useful sequence information across various bioinformatics tasks (Bi et al., 2022; Chen et al., 2023; Wang et al., 2024; Zeng et al., 2023). Additionally, their capacity to handle sequences of varying lengths enables the Allocator framework to operate without constraints on sequence length. Specifically, the *k*-mer encoding calculates the frequency of *k* adjacent nucleic acids. In this study, we considered the frequencies of 1-mer, 2-mer, 3-mer, 4-mer and 5-mer, resulting in a feature vector with a dimensionality of 1364. On the other hand, CKSNAP represents the frequency of nucleic acid pairs by any *k* nucleic acid. We used *k* values of 0, 1, 2, 3, 4 and 5, leading to a 96-dimensional feature vector. The feature extraction process described above was conducted using the iLearn toolkit and detailed descriptions of these encoding methods can be found (Chen et al., 2020).

Incorporating the RNA’s secondary structure into the model can offer valuable insights into its folding patterns, including structural elements like stem loops and pseudoknots. However, experimentally determined RNA secondary structures are minimal and insufficient for our use. Therefore, we employed the predictive secondary structures to extract graph structural features. For each RNA secondary structure, we represent it as a graph with individual bases as nodes. The graph is constructed using two types of edges: one connecting adjacent bases and the other representing base-pairing interactions derived from a predictive tool. We have prepared two distinct graphs predicted by two popular prediction tools, *i*.*e*., RNAFold (Lorenz et al., 2011) and LinearFold (Huang et al., 2019), to diversify the resources of predicted structures and enhance the reliability of structural features extracted. We used the dot-bracket notation of mRNAs generated by LinearFold and RNAFold to construct graphs. Dot-bracket notation is a commonly used representation of RNA secondary structure that uses “(” and “)” to indicate pairing between bases and “.” to denote unpaired bases. **Algorithm 1** shows the details of building RNA graphs using dot-bracket annotation inputs. The input is the dot-bracket notation string **D**, and the output is the graph object **G**. Variable **bases** is a list used to store the indexes of the paired bases. Subsequently, a **for** loop iterates through the dot-bracket notation string, representing each base as a node of **G**, adding edges of type “base pair” between nodes with complementary bases, and adding edges of type “adjacent” between neighbouring nodes. Finally, the algorithm returns the graph object **G**, which is used as an input to the graph neural network to learn the structural features of the mRNA.

Both these two types of graphs include the formation of hydrogen bonds and phosphodiester bonds. In addition, each node is denoted by a 10-dimensional feature vector that integrates four different encodings: one-hot, NCP (nucleotide chemical property), EIIP (electronion interaction pseudopotentials), and ANF (accumulated nucleotide frequency) (Liu et al., 2021). (More detailed descriptions of the node features are provided in **Supplementary Table S1**).

### Sequence context feature extractor

To facilitate the feature extractor’s capabilities at both local and global levels, we introduced multiple types of neural network architectures into our sequence context feature extractors. These two sequence context feature extractors share a standard set of components, which primarily include an MLP layer and a multi-head self-attention layer. As a core extractor component, the multi-head self-attention enables models to focus on different aspects of important information

#### Algorithm 1: Dot-bracket to graph

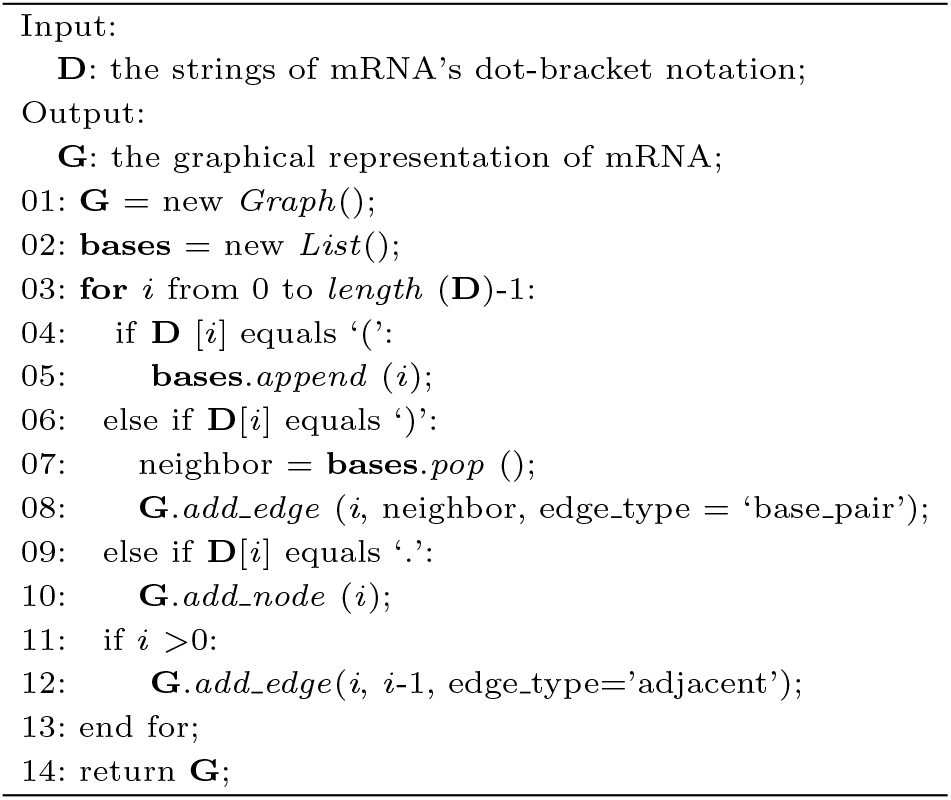

from multiple subspaces and weigh the importance of individual words within the input sequences, thus improving the quality of the feature representations learned by the framework. This mechanism is achieved by combining multiple attention heads, each relying on the scaled dot products to calculate the attention scores. The specific process can be defined as follows:

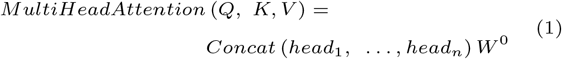

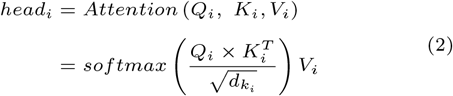

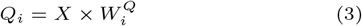

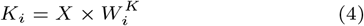

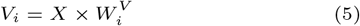

where *X* represents the input of attention module, 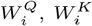, 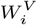 are learnable parameter matrices, and 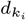 is the dimension of matrix *K*_*i*_.

### Graph structural feature extractor

In this study, we employed a graph isomorphic network (GIN) to serve as the Allocator’s graph structural feature extractors as it performed better than other GNN variants in our preliminary tests. Graph Isomorphic Network (GIN) (Xu et al., 2019) is designed to aggregate information from neighbouring nodes, as formulated below:

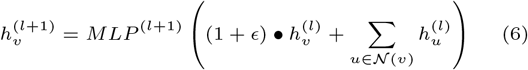

where 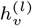 is the representation of node *v* at layer *l, ϵ* is a learnable parameter, and *MLP* ^(*l*+1)^ is a multilayer perceptron used to update the node’s representations. Following the GIN layer, we employed a set2set layer, acting as a global graph pooling mechanism, to capture and summarize information from the entire set. This layer outputs a fixed-size representation for subsequent combinations with other extractors’ outputs. To simplify the code writing, we used the PyTorch Geometric package (Fey & Lenssen, 2019) to implement the construction of a graph neural network. For more detailed model parameters for the Allocator, please refer to **Supplementary Table S2**.

### Prediction module

Allocator’s prediction module is a feedforward network with two layers. After concatenating the features learned by these four feature extractors, the prediction module performs the linear transformation and then employs a sigmoid function in the second layer of the prediction module to produce the final output, which is formulated as follows:

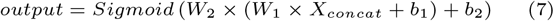

where *W*_1_, *W*_2_, *b*_1_, and *b*_2_ are learnable parameters.

### Model training and evaluation

During the training of Allocator, we employed the Adam optimization approach (Kingma & Ba, 2017) to enhance the training process and optimize the model’s parameters. To measure the dissimilarity between the true labels and the predicted probabilities, we used the binary cross-entropy function that is defined as follows:

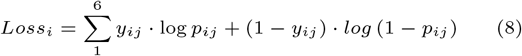

where *y*_*ij*_ takes the value 0 or 1, and *p*_*ij*_ takes the value in [0, 1]. When the *y*_*ij*_ value is 0, it means that the mRNA sample *x*_*i*_ does not belong to the corresponding mRNA subcellular localization compartment *j* (1, 2, 3, 4, 5, 6); on the contrary, if the *y*_*ij*_ value is 1, it means that the mRNA sample *x*_*i*_ is located in the corresponding compartment *j*. Our training objective is to make the predicted probability value as close to 1 or 0 as possible, with the model providing separate predicted probability values for each location compartment.

To evaluate Allocator’s overall performance, we employed six performance evaluation metrics designed explicitly for multi-label problems (Bi et al., 2022; Gopal & Yang, 2010), including example-based accuracy (*Acc*_*exam*_), average precision, coverage, one-error, ranking loss, and hamming loss. These metrics offer a comprehensive assessment of Allocator’s capabilities across all six labels/locations. The descriptions of these metrics are provided in the “Performance evaluation metrics” section in the **Supplementary Material**.

## Results and discussion

### Allocator’s multi-label performance

DM3Loc used one-hot features to achieve competitive performance; however, it might overlook capturing relationships between nucleotide sequences and structural information. The Allocator’s key innovation lies in the extraction of the RNA secondary structural features using graph deep learning techniques. We sought to investigate whether these graph structural features can provide additional insights. As part of our exploration to explore the effect of RNA secondary structure, we retrained a new model only utilizing sequence context extractors with sequence features including k-mer and CKSNAP, that is, Allocator (seq). This model differs from the original Allocator (seq + struct). Both models were trained with a learning rate of 0.0003 and a dropout rate of 0.1, lasting for 200 epochs. We conducted the performance comparison among Allocator (seq), Allocator (seq + struct), and DM3Loc, and the results are illustrated in **Figure 2**. The results demonstrated that Allocator (seq + struct) leads to a notable improvement in terms of (*Acc*_*exam*_). Additionally, it reduces the coverage, ranking loss, and hamming loss when compared to Allocator (seq). This demonstrates that integrating the secondary structure clearly boosts predictive performance, highlighting the value of using multi-view features.

**Fig. 2.**
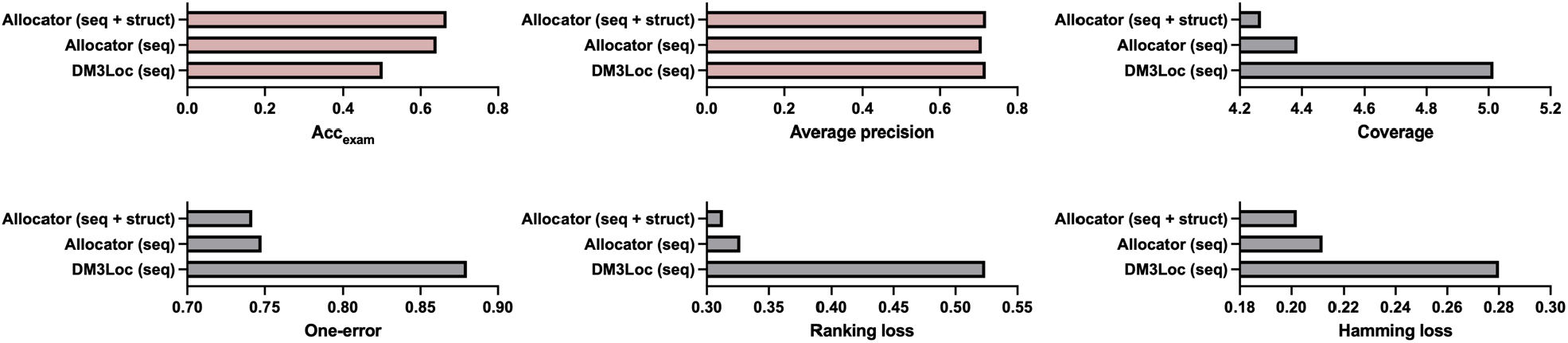
The performance of Allocator across six multi-label evaluation metrics.

In addition to the multi-label evaluation, we also conducted a single-label assessment of the Allocator’s performance. We used accuracy and Matthews correlation coefficient (MCC) to measure the efficacy of each label/localization (**See supplementary Table S3**). Allocator’s accuracies are higher than DM3Loc in all six localization predictions. Notably, for the exosome’s prediction, the Allocator model achieves an accuracy of 98.80%, surpassing DM3Loc’s 73.71% by approximately 25 percentage points. However, Allocator’s MCC performance falls short of that achieved by DM3Loc. We found that the Allocator’s performance is not satisfactory for imbalanced labels in the case of exosome, membrane and ER localization. This can be attributed to the model’s training direction, which emphasizes label relationships and tends to favour the majority classes. In contrast, DM3Loc defined a range of cutoffs for each localization, leading to an increase in MCC but a decrease in accuracy. Indeed, addressing this inherent problem requires the inclusion of more training samples in the future. Overall, while Allocator may not excel in terms of MCC, it still offers a valuable perspective for uncovering label relationships.

### Architecture analysis

The feature extractor module serves as the essential core of the Allocator architecture and is responsible for extracting intricate patterns from multi-view inputs of mRNAs, including primary sequences and secondary structures. To understand the contributions of each component in Allocator, we conducted experimental analyses from the views of sequence context feature extractors, graph structural feature extractors, and the entire architecture.

### Sequence context feature extractor study

We explored four network combinations to optimize the sequence context feature extractor, which includes using only MLP, MLP+attention, CNN+LSTM (Liu et al., 2022), and CNN+LSTM+attention (Li et al., 2023). The MLP consists of three hidden layers, while both CNN and LSTM only have a single layer. As illustrated in **Table 2**, the combination of MLP+attention achieves the best results in terms of Accexam and average precision, along with yielding the lowest coverage, one-error, and hamming loss. The combination of CNN and LSTM (CNN+LSTM) did not demonstrate superior performance, possibly due to the input features not being context-independent. Furthermore, we observed a significant improvement when the attention mechanism was added compared to the model without attention. Therefore, we adopted the combination of MLP and attention (MLP+attention) for the sequence context feature extractors, aiming to extract in-depth attributes from sequence-based features.

**Table 2.** Performance comparison on the test set using different neural network types.

| Type | Accexam | Average precision | Coverage | One-error | Ranking loss | Hamming loss |
| --- | --- | --- | --- | --- | --- | --- |
| S: MLP +attention & G: GIN (Allocator) | 0.667 | 0.719 | 4.268 | 0.742 | 0.313 | 0.202 |
| S: MLP | 0.647 | 0.709 | 4.344 | 0.770 | 0.337 | 0.218 |
| S: CNN + LSTM | 0.638 | 0.716 | 4.421 | 0.763 | 0.355 | 0.221 |
| S: CNN + LSTM + attention | 0.650 | 0.708 | 4.388 | 0.762 | 0.312 | 0.205 |
| G: GAT | 0.650 | 0.710 | 4.347 | 0.748 | 0.323 | 0.209 |
| G: GCN | 0.635 | 0.716 | 4.439 | 0.744 | 0.352 | 0.216 |
| G: MPNN | 0.640 | 0.707 | 4.355 | 0.760 | 0.346 | 0.221 |
| Without S-kmer | 0.656 | 0.714 | 4.360 | 0.753 | 0.309 | 0.201 |
| Without S-CKSNAP | 0.646 | 0.712 | 4.352 | 0.763 | 0.347 | 0.219 |
| Without G-LinearFold-SS | 0.652 | 0.716 | 4.334 | 0.758 | 0.333 | 0.212 |
| Without G-RNAFold-SS | 0.641 | 0.708 | 4.369 | 0.776 | 0.347 | 0.222 |

\*Abberiations: sequence context feature extractor (S); graph structural feature extractor (G).

### Graph structural feature extractor study

As mentioned in the introduction section, GIN, GAT, GCN, and MPNN are different types of graph neural networks, each possessing unique characteristics and approaches to handling graph-structural data. To guide the development of our graph structural feature extractors, we explored these four network types to evaluate their impact on model performance. The results suggest that the GIN-based model consistently attains the highest levels of *Acc*_*exam*_ and average precision, thereby establishing GIN as the optimal choice for Allocator’s graph structural feature extractors. The application of graph neural networks to identify mRNA subcellular localization can be regarded as a graph classification task, and the graph representation capability of a graph neural network determines the classification capability of the prediction model. One reason why GIN has been shown to outperform other GNN variants such as GCN in aggregating and updating node features is that it introduces MLPs to aggregate and update node features (Xu et al., 2019). Graph neural networks represent graphs by aggregating node features, and commonly employed aggregators, such as maximum or mean, make graph neural networks unable to distinguish between very simple graph structures, reducing the network’s ability to characterize graphs. In contrast, MLPs can fit any function approximately, which gives GIN a stronger graph representation capability. On the other hand, in biological networks such as protein structures, RNA structures, and gene networks, protein RNA or gene structures with similar structures have similar functional properties. the graph of mRNA secondary structure contains only four types of nodes, which makes the converted graphs have substructures with high similarity. Therefore, GIN, which has an advantage in distinguishing graph structures, provide stronger graph representation learning ability than other graph neural networks such as GCN in this study.

### Ablation study

After determining the multi-view feature extractors of Allocator, we aimed to test whether this framework is overly complex and redundant. We performed ablation studies to evaluate the impact of removing each extractor individually. These studies involved eliminating the sequence context feature extractor for *k*-mer (*i*.*e*., S-kmer), the removal of the sequence context feature extractor for CKSNAP (S-CKSNAP), the exclusion of the graph structural feature extractor for secondary structure from Linearfold (G-linearFold-SS), and the exclusion of the graph structural feature extractor for secondary structure from RNAfold (G-RNAFold-SS). As indicated in **Table 2**, the removal of any single feature extractor led to a decrease in *Acc*_*exam*_ and average precision compared to the full Allocator setup. Notably, in contrast to S-kmer and G-LinearFold-SS, S-CKSNAP and G-RNAfold-SS exhibit a more pronounced influence on Allocator. As a result, we decided to retain all four extractors to maintain a more robust model capacity.

### Analysis of single-localization

In this section, we explored the impact of the Allocator’s predictions on single localizations. Firstly, we performed a statistical analysis using the test set, comparing the actual and predicted samples of six different localizations. As depicted in **Figure 3A**, we observed that the distributions for exosome and ribosome closely matched between the actual and predicted samples. In contrast, for the nucleus and cytosol, the predicted samples exceeded the actual samples, while for membrane and ER, the opposite was observed. One potential explanation is the impact of imbalanced samples across different localizations, as these imbalances appear to have a cross-influence on one another. Nevertheless, it’s crucial to highlight that, on the whole, the actual labels and those predicted remain highly similar, providing evidence of the Allocator’s outstanding predictive ability.

**Fig. 3.**
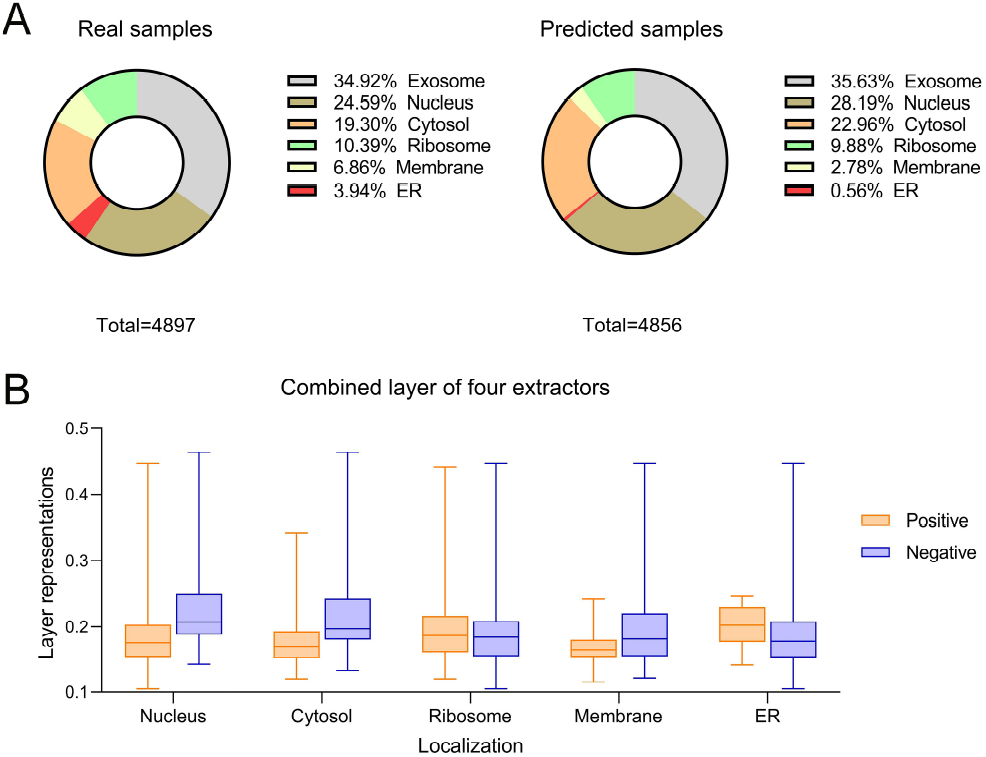
(A) shows the distribution of actual and predicted samples for all six localizations; (B) displays the layer output representations for positives and negatives in five localizations, excluding exosome.

Afterwards, we selected correctly predicted samples for each localization separately and observed the layer representations for each of the four extractor combinations. The reason for not being able to choose the same samples is that their feature outputs are the same since this is a multi-label problem, and thus, no differences can be observed. Furthermore, we excluded the exosome category from our analysis due to its extreme imbalance, and we could not obtain a sufficient number of negatives for this category. **Figure 3B** displays the average representation values for positives and negatives across the remaining five locations. Notably, we observe significant differences in the categories of nucleus, cytosol, membrane and ER. This highlights the effectiveness of combining the outputs from the sequence context feature and graph structural feature extractors, thereby reinforcing the superiority of Allocator.

### Model interpretation

To investigate the key substructures critical for localization, we employed the model interpretation algorithm GNNExplainer (Ying et al., 2019), designed specifically for elucidating graph neural network models to interpret the Allocator model. The node feature importance evaluation results shown in **Figure 4(A)** reveal that ANF and EIIP are the most significant features for all six localizations. In contrast, NCP and one-hot encoding were less significant, likely due to their binary nature and lower information density compared to ANF and EIIP’s continuous attributes. We then identified eleven key substructures and analyzed their distributions across the six localizations. Specifically, from the top 300 edges based on important scores, we noticed that some adjacent/paired edges form a partial substructure, with at least 3 bases involved. We identified a subset of substructures and conducted a detailed investigation into their proportions across each localization (*N*_*sub*_*/N*_*total*_ in each localization), as shown in **Figure 4(B)**. Within these substructures, A==U is more prevalent than G==C, indicating its possibly vital function. Notably, we found a unique substructure in membrane localization: the A*↔*A==A substructure, suggesting a potential link to membrane localization.

**Fig. 4.**
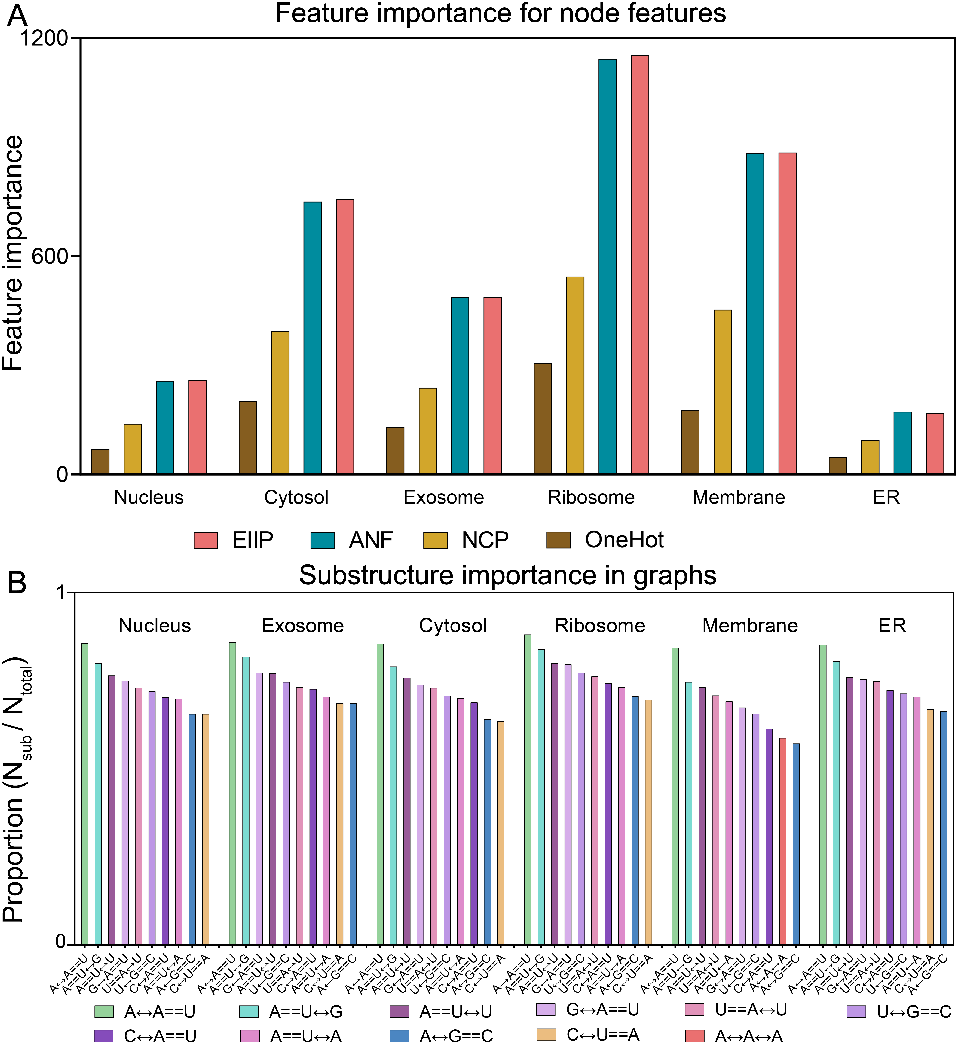
(A) shows the importance of four features for six localization categories; (B) displays the first ten important substructures for six localizations. Where == denotes base pairing and *↔* indicates neighboring

To more explicitly demonstrate the effectiveness of secondary structure features in enhancing Allocator’s feature representation learning ability, we selected four case study mRNAs and used their GNNExplainer results for visualization. We aim to illustrate how secondary structures can enhance the feature representations of mRNAs, thereby improving prediction accuracy. The selected mRNAs include STUB1 (Gene name: STUB1, localized to Nucleus, Exosome, Cytosol, Ribosome, Membrane, and ER); HIST4H4 (Gene name: HIST4H4, localized to Nucleus, Exosome, and Ribosome); TMEM238 (Gene name: TMEM238, localized to Exosome and Ribosome;) and SPINK9 (Gene name: SPINK9, localized to Exosome). For each mRNA, we used GNNExplainer to calculate the importance scores for each edge in its graph. We selected the edges ranked in the top three hundred by importance to identify key subgraphs from the graph. We attempted to identify important subgraphs by selecting edges with the highest importance scores. Ideally, these subgraphs are composed of structural connections of multiple bases and the results are visualized in Figure 5. The heatmaps in Figure 5 (i, iii, v, and vii) visualize the edge importance scores within identified important subgraphs of four case study mRNAs. The heatmaps’ horizontal and vertical coordinates correspond to the subgraph nodes and their respective positions within the graph. The secondary structure visualisations of important subgraphs are presented in Figures 5 (ii, iv, vi, and viii) and the full structure visualisations generated by RNAfold are illustrated in Supplementary Figures S1-S4. The four different types of bases (A, G, C, U) are represented by different coloured circles and labelled with position information. The two types of edges are indicated by grey lines for adjacent edges and red lines for paired edges.

**Fig. 5.**
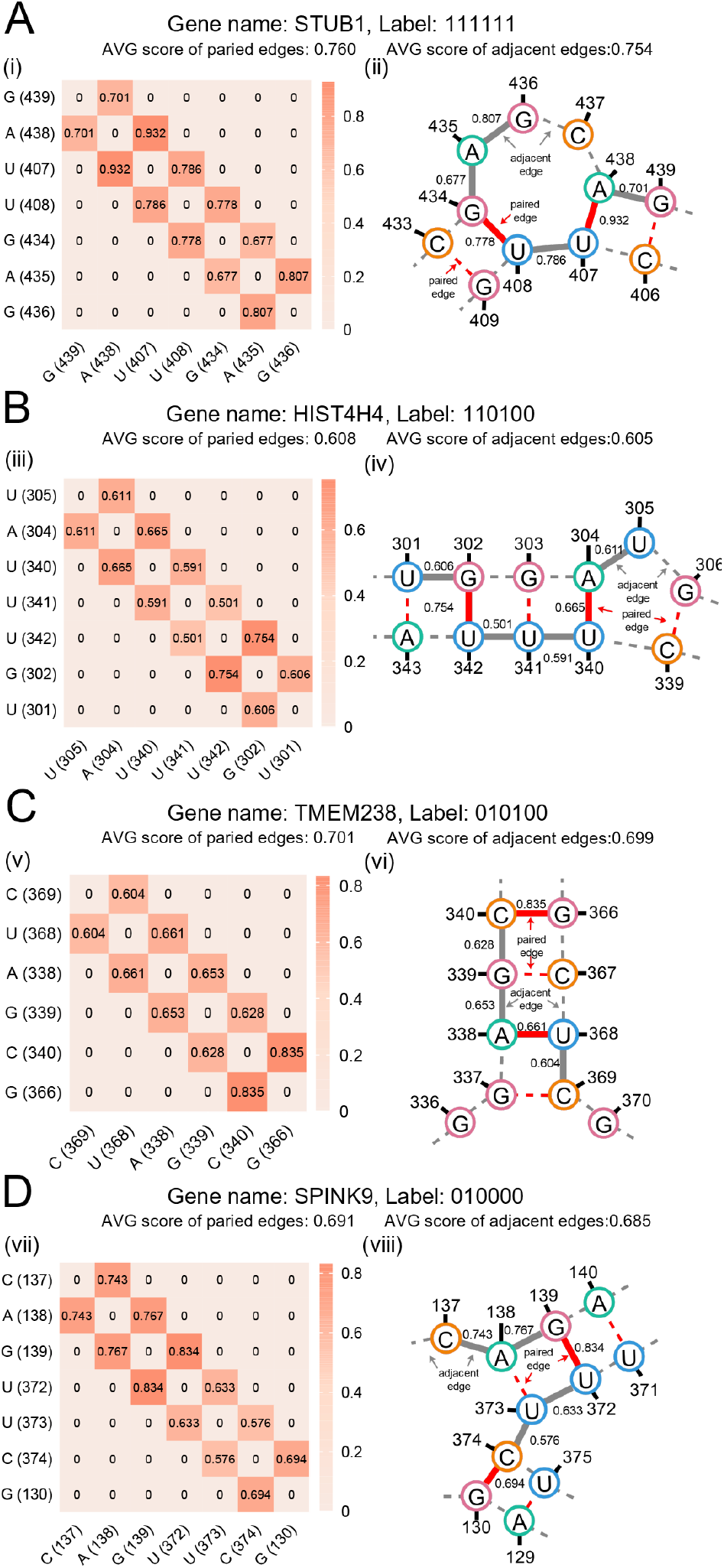
Visualisation of the Secondary structure and heat map of four case study mRNAs. (A) STUB1: localized to Nucleus, Exosome, Cytosol, Ribosome, Membrane, and ER; (B) HIST4H4: localized to Nucleus, Exosome, and Ribosome; (C) TMEM238: localized to Exosome and Ribosome; and (D) SPINK9: localized to Exosome.

We found that the importance scores of some paired edges are larger than those of adjacent edges in these subgraphs and we statistically concluded that the average importance score of paired edges is greater than that of adjacent edges for all four case study mRNAs. Specifically, in STUB1, the identified subgraph “GAUUGAG” is located on a loop of STUB1 (Figure 5i and 5ii), and the importance score of the paired edge between A (438) and U (407) is much larger than that of the adjacent edge between A (438) and G (439) (0.932*>*0.701), and the importance scores of the paired edges between G (434) and U (408) are larger than those of the adjacent edges between G (434) and A (435) (0.778*>*0.677), and slightly smaller than those of the adjacent edges between U (408) and U (407) (0.778*<*0.786). In the subgraph “UAUUUGU” found on the stem-loop (Figure 5iii and 5iv) of HIST4H4, the importance score of the paired edge between G (302) and U (342) is larger than that of the adjacent edge between G (302) and U (301) (0.754*>*0.606), and also larger than that of the adjacent edge between U (342) and U (341) (0.754*>*0.501), and the importance score of paired edges between A (304) and U (340) are larger than those of the adjacent edges between A (304) and U (305) (0.665*>*0.611), and also larger than that of the adjacent edge between U (340) and U (341) (0.665*>*0.591). Similarly, in mRNA SPINK9 (Figure 5vii and 5viii), the subgraph “CAGUUCG” on the stem-loop was found to have two paired edges between G (139) and U (372), and between C (374) and G (130), with scores greater than those of the adjacent edges adjacent to them (0.834*>*0.767, 0.834*>*0.633, 0.694*>*0.576). In mRNA TMEM238 (Figure 5v and 5vi), the pairwise edges of the subgraph “CUAGCG” located on the stem are the edges between C (340) and G (366), and between U (368) and A (338), which also score higher than their adjacent edges (0.835*>*0.628, 0.661*>*0.604, 0.661*>*0.653). In summary, we infer that Allocator benefits from the ability to capture important secondary structure information.

### Webserver and software development

To facilitate easy access to Allocator, we have developed a user-friendly web server, which is freely available at http://allocator.unimelb-biotools.cloud.edu.au. The web server is hosted on a robust Linux server, equipped with four cores, 16 GB of memory, and a spacious 200 GB hard disk. This powerful configuration enables the server to handle various tasks and deliver a smooth user experience. With Allocators’s web server, users are required to upload or paste the mRNA sequences in FASTA format to submit an online prediction task. Once the prediction is completed, the results will be presented in a tabular format. Users can download the results in different file formats, such as EXCEL, TXT, and CSV. We also offer a detailed guide on the ‘Help’ page of our website, explaining the specific steps for using Allocator. Please note that the online prediction can simultaneously accommodate up to 5 mRNA sequences. For prediction involving a larger range of sequences, please download the standalone software of Allocator for fast and efficient local predictions.

## Discussion and conclusion

The widespread identification of mRNA subcellular localizations is crucial in understanding intricate gene regulatory mechanisms. To complement labour-intensive wet lab experiments, several computational approaches have been proposed for predicting mRNA subcellular localizations. However, most existing methods overlook the multi-localization aspect of mRNA and treat it as a simple single-label multi-class task. While a multi-label and multi-class model, DM3Loc, has been developed based on the self-attention neural network, there is a gap in addressing the complexities of mRNA subcellular localizations due to the dynamic cellular processes. The secondary structure of mRNA is closely related to its functionality and may contain crucial information regarding transport, such as *cis*-localization element. Building on this insight, we introduced a deep learning model named Allocator, designed to identify multiple mRNA localizations simultaneously. Allocator not only considers mRNA primary sequence to generate sequence context features but also raises the secondary structure to create the graph representations. The main components of Allocator models consist of two sequence context feature extractors specialized for sequence features and two graph structural feature extractors for secondary structural representations. This heterogeneous network architecture empowers the Allocator models to exhibit robust performance. In summary, this paper showcases the effectiveness of the Allocator framework, which holds promise for application in various biological analysis tasks in future research.

Despite achieving satisfactory results in revealing label relationships, Allocator still faces challenges performing well on imbalanced labels. Additionally, it is essential to emphasize that the mRNA secondary structure used in this work is derived from prediction tools and may not accurately represent the actual secondary structure of mRNA, potentially impacting the model’s performance. We believe that with the ongoing advancement and expansion of experimental data and databases, these challenges can be effectively mitigated and resolved, leading to an enhancement in the model’s capabilities.

## Acknowledgments

Fuyi Li and Yue Bi contributed equally to this work.

## Author contributions

FL and SP conceived the study; FL, YB, XG, XT, and CW developed the method, performed experiments, and analyzed results; FL and SP wrote the manuscript.

## Data and code availability

The datasets and source codes are publicly available at https://github.com/lifuyi774/Allocator.

