## Supplementary Material for "Advancing mRNA subcellular localization prediction with graph neural network and RNA structure"

### 1 Supplementary Tables

Table S1: The encoding schemes of NCP (nucleotide chemical property), EIIP (electronion interaction pseudopotentials) and ANF (accumulated nucleotide frequency).

| | Nucleotide types | NCP | EIIP | ANF |
|---|---|---|---|---|
| A | [0,1,0,0,0] | [1,1,1] | 0.1260 | d |
| G | [0,0,1,0,0] | [1,0,0] | 0.0806 | d |
| C | [0,0,0,1,0] | [0,1,0] | 0.1340 | d |
| T/U | [0,0,0,0,1] | [0,0,1] | 0.1335 | d |

Table S2: Allocator's parameter setting.

| Parameter | Setting |
|---|---|
| Number of GIN input features | 10 |
| Number of MLPs input features | 96/1364 |
| Hidden layer dimension | 64 |
| Number of output classes | 6 |
| Dropout rate | 0.1 |
| Activate function | Relu |
| Optimization algorithm | Adam (learning rate:0.0003) |
| Batch size | 128 |
| Batch normalization | True |

Table S3: Performance for each label/localization between DM3Loc and allocator.

| Localization | MCC | | Accuracy | |
|---|---|---|---|---|
| | DM3Loc | Allocator | DM3Loc | Allocator |
| Nucleus | 0.3483 | 0.3309 | 0.6817 | 0.7362 |
| Exosome | 0.0422 | -0.0026 | 0.7371 | 0.9880 |
| Cytosol | 0.1859 | 0.3014 | 0.5783 | 0.6563 |
| Ribosome | 0.3531 | 0.3054 | 0.6811 | 0.7145 |
| Membrane | 0.2999 | 0.2292 | 0.7805 | 0.8083 |
| ER | 0.0241 | 0.1393 | 0.8614 | 0.8864 |

### 2 Supplementary Material

#### 2.1 ANF description

The ANF encoding contains the information and distribution of each nucleotide in the RNA sequence, and the calculation formula is as follows:

$$d_i = \frac{1}{|s_i|}\sum_{j=1}^{l} f\left(s_i\right),\ f(q) = \begin{cases} 1\ if\ s_i = 1, \\ 0\ other\ case \end{cases} \tag{1}$$

where $l$ represents the length of the mRNA sequence, and $s_i$ is the length of the $i$-th prefix sequence fragment$\{s_1,\ s_2,\ldots,s_i\ \}$ in the mRNA sequence, $q\ \epsilon\ \{\text{A,C,G,U}\}$. For instance, when considering the sequence 'UCGGUCAUCG', the ANF encoding values for 'U' at positions 1, 5, and 8 in the sequence are 1 (1/1), 0.4 (2/5), and 0.375 (3/8), respectively.

#### 2.2 Performance evaluation metrics

To evaluate Allocator's overall performance, we utilized six evaluation metrics designed explicitly for multi-label problems, including example-based accuracy ($Acc_{exam}$), average precision, coverage, one-error, ranking loss, and hamming loss. These metrics offer a comprehensive assessment of Allocator's capabilities across all six labels/locations. The descriptions of these metrics are provided below:

$$Acc_{exam} = \frac{1}{t}\sum_{i=1}^{t}\frac{|P_i \cap Y_i|}{|P_i \cup Y_i|} \tag{2}$$

$$\begin{aligned} Average\ Precision = \frac{1}{t}\sum_{i=1}^{t}\frac{1}{|Y_i|} \\ \sum_{y\in Y_i}\frac{|\{y'\,|Rank_f\,(x_i,y') \leq Rank_f\,(x_i,y)\,, y' \in Y_i\,\}|}{Rank_f\,(x_i,y)} \end{aligned} \tag{3}$$

$$Coverage = \frac{1}{t}\sum_{i=1}^{t} max_{y'\in Y_i} Rank\,[f\,(x_i,y')] - 1 \tag{4}$$

$$One - error = \frac{1}{t}\sum_{i=1}^{t} I\left(\arg max_{y'\in Y_i} f\,(x_i,y') \notin Y_i\right) \tag{5}$$

$$\begin{aligned} Ranking\ Loss = \frac{1}{t}\sum_{i=1}^{t} \\ \frac{1}{|Y_i|\,\left|\bar{Y}_i\right|} I\left(f\,(x_i,y') \leq f\,(x_i,y'')\,, y' \in Y_i,\ y'' \in \bar{Y}_i\right) \end{aligned} \tag{6}$$

$$Hamming\ Loss = \frac{1}{t}\sum_{i=1}^{t}\frac{1}{q}\,|P_i \Delta Y_i| \tag{7}$$

where $f(\cdot)$ represents the classifier; $Y_i$ and $\overline{Y}_i$ represent the set of real labels and the complement of this set, respectively; $P_i$ represents the set of predicted labels; $(x_i, y_i)$ $\{1 \leq i \leq t\}$ represents a multi-label instance; $Rank_f(x, y)$ indicates the descending ranking of $y$ in $Y$; $|\cdot|$ indicates the cardinality of the set, $q = |Y_i|$; $\varDelta$ is the symmetric difference between two sets; and $I(\cdot)$ indicates the count that satisfies the condition.

### 3 Supplementary Figures

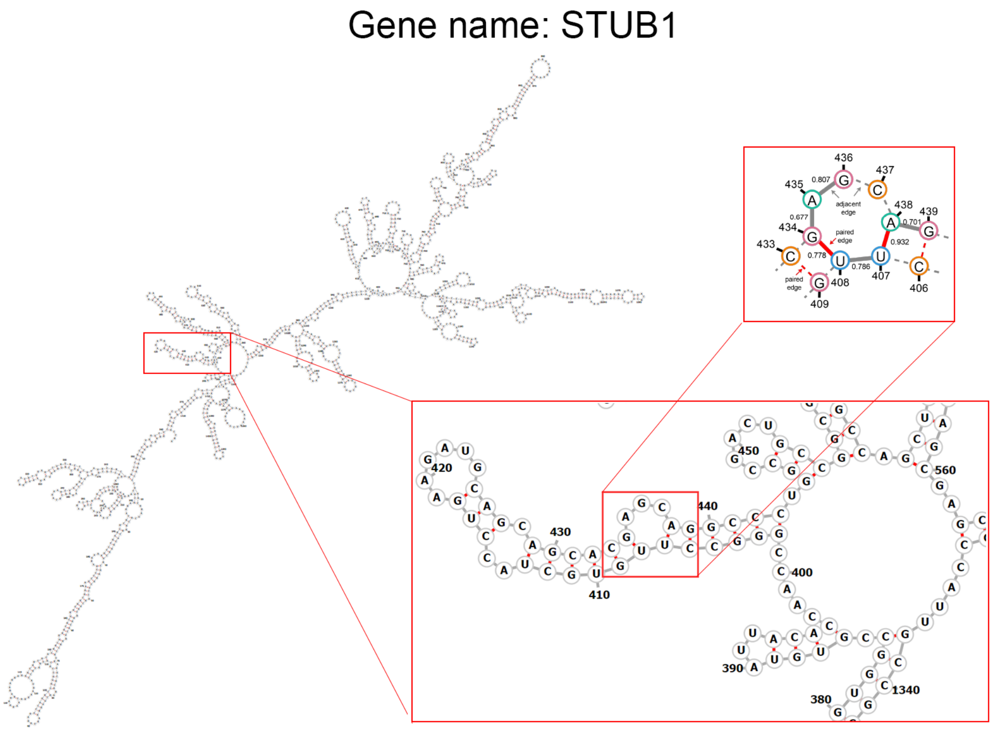


**Fig. S1.** Schematic representation of the secondary structure (predicted by RNAFold) of mRNA STUB1.

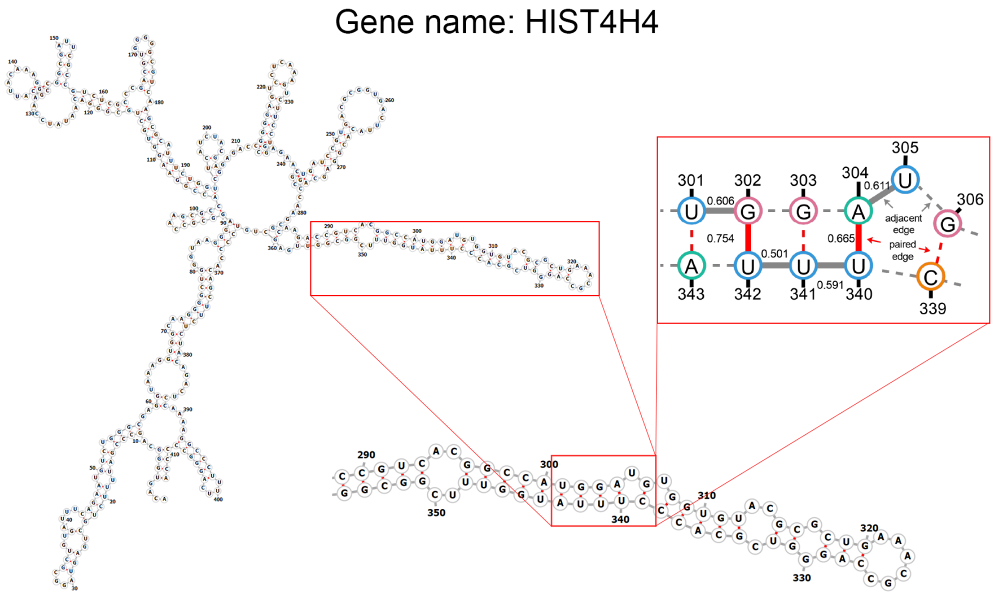


**Fig. S2.** Schematic representation of the secondary structure (predicted by RNAFold) of mRNA HIST4H4.

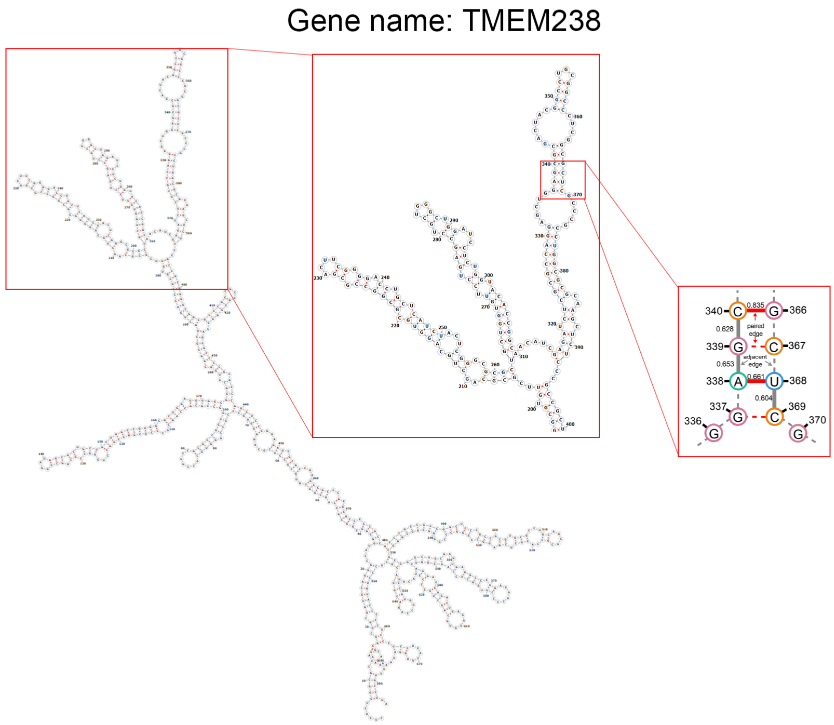


**Fig. S3.** Schematic representation of the secondary structure (predicted by RNAFold) of mRNA TMEM238.

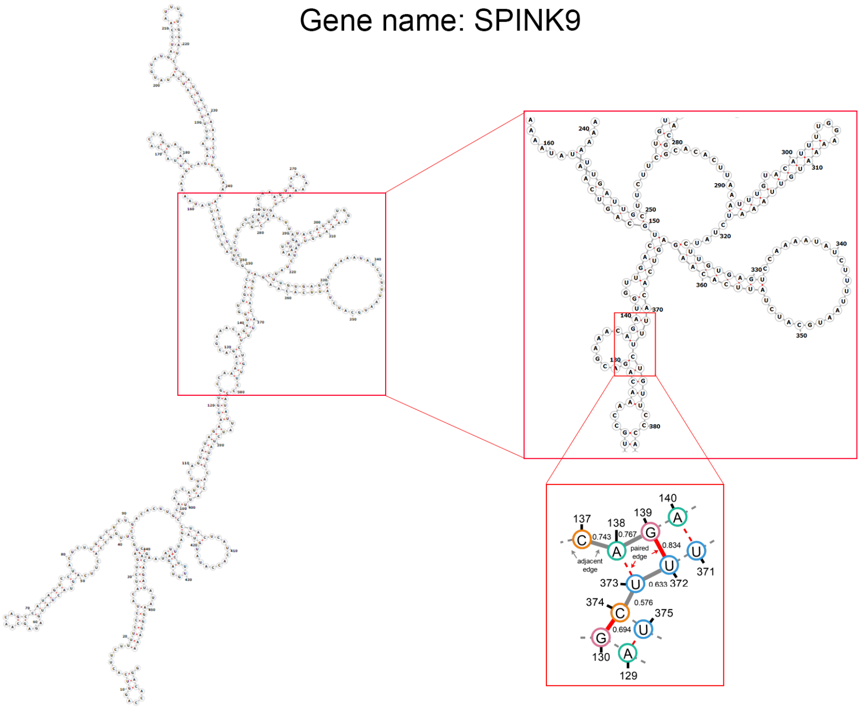


**Fig. S4.** Schematic representation of the secondary structure (predicted by RNAFold) of mRNA SPINK9.
